# Ellagic acid: a potential inhibitor of enhancer of zeste homolog-2 and protein arginine methyltransferase-5

**DOI:** 10.1101/2024.05.22.595443

**Authors:** Kirankumar Nalla, Biji Chatterjee, Jagadeesha Poyya, Aishwarya Swain, Krishna Ghosh, Archana Pan, Chandrashekhar G Joshi, Bramanandam Manavathi, Santosh R Kanade

## Abstract

Dysregulation of epigenetic processes, characterized by aberrant DNA methylation patterns and histone modifications, is a hallmark of cancer, driving its initiation, progression, and metastasis by silencing tumor suppressor genes or activating oncogenes. Perturbations in histone modifications such as H3K27me3 by EZH2 and H4R3me2s by PRMT5 play significant roles in these epigenetic alterations, disrupting normal gene expression and facilitating oncogene activation while suppressing tumor suppressor genes. Consequently, inhibitors targeting enzymes involved in DNA methylation, histone modification, or chromatin remodeling, such as PRMTs and PRC complexes, are promising anti-cancer agents, with several undergoing pre-clinical and clinical trials. Our screening of a phytochemical library revealed ellagic acid as an effective inhibitor of both EZH2 and PRMT5. Ellagic acid interacts strongly with the EZH2 and PRMT5:MEP50 complex, binding to their active sites through π-cation interactions and hydrogen bonds. Surface Plasmon Resonance study confirmed potent binding affinities of ellagic acid, with KD values of 3.28E-06 and 6.54E-05 for EZH2 and PRMT5:MEP50 respectively. *In-vitro* assays validated inhibitory effects on EZH2 and PRMT5:MEP50 by reducing the levels of their catalytic products H3K27me3 and H4R3me2s respectively and induction of autophagy and apoptosis. Further *In-vivo* studies using mouse xenografts further demonstrated significant tumor size reductions upon oral administration of ellagic acid, with decreased expression of the proliferative marker ki67 and histone repressive marks. Taken together we showed that inhibition of EZH2 and PRMT5:MEP50 by ellagic acid could be used to develop breast cancer therapeutic drug.

## Introduction

Cancer, a leading cause of global mortality, continues to pose significant challenges to healthcare systems worldwide. With millions of new cases diagnosed each year, the burden of cancer is projected to escalate substantially in the coming decades ^(1, 2).^ As per WHO latest reports, cancer stands as a significant factor in worldwide mortality, responsible for approximately one-sixth of all deaths and impacting nearly every household ^(1).^ In 2022 alone, there were an estimated 20 million fresh instances of cancer, leading to 9.7 million fatalities globally. Projections suggest that by 2050, the burden of cancer will surge by approximately 77%, placing additional strain on health systems, individuals, and communities ^(2).^ In 2023, in the United States approximately 2.0 million people received a cancer diagnosis, with breast cancer being the most common among women and prostate cancer among men ^(3).^ Breast and prostate cancer may be influenced by factors like physical activity, obesity, age, family history, and oral contraceptive use. Among the diverse factors influencing cancer development and progression, epigenetic dysregulation stands out as a hallmark feature ^(4).^ Its progression involves genetic and epigenetic changes, particularly through DNA modification like methylation and histone modifications like acetylation methylation, ubiquitination, phosphorylation, and sumoylation. These alterations affect structure of chromatin and various pathways linked to breast and prostate cancer development ^(4).^

Epigenetic dysregulation profoundly influences cancer initiation, progression, and metastasis. This dysregulation primarily manifests through aberrant DNA methylation patterns and alterations in histone modifications, which are critical in gene expression and chromatin structure. Uncommon DNA methylation can result in the tumor suppressor genes silencing, essential guardians of cellular integrity and regulators of cell growth, division, and apoptosis. Conversely, it can lead to the activation of oncogenes, with cell proliferation and survival enhancement, contributing to malignant transformation ^(5, 6).^ Similarly, perturbations in histone modifications, such as H3K27me3 catalyzed by EZH2(7) and H4R3me2s mediated by PRMT5(8), play pivotal roles in driving these epigenetic alterations in cancer cells. Histone modifications regulate the accessibility of chromatin and modulate gene expression patterns. When dysregulated, they can disrupt the delicate balance between active and repressive chromatin states, thereby promoting aberrant gene expression profile conducive to tumorigenesis ^(9).^ For instance, EZH2-mediated H3K27me3 trimethylation typically leads to transcriptional repression, effectively silencing tumor suppressor genes and promoting oncogene activation ^(10).^ Conversely, PRMT5-mediated H4R3me2s symmetric dimethylation may also contribute to transcriptional repression, further exacerbating the dysregulation of gene expression networks involved in cancer progression ^(8, 9)^ Understanding these epigenetic alterations is crucial for unraveling the molecular mechanisms underpinning cancer development and identifying potential therapeutic targets to restore normal epigenetic regulation.

Despite their distinct roles, EZH2 and PRMT5 share a commonality as histone methyltransferases. These enzymes selectively and specifically catalyze the methylation of lysine amino acid or arginine amino acid residues in target histone and non-histone proteins, thereby modulating chromatin structure and gene expression patterns. This classification places EZH2 and PRMT5 in the broader categories of lysine methyltransferases and arginine methyltransferases, respectively. Their actions contribute to the epigenetic regulation of gene expression, impacting various cellular processes and ultimately influencing cancer development and progression. PRMTs are responsible for arginine methylation on histone and non-histones, classified into type I, type II, and type III. Type I catalyze asymmetric dimethylation of arginine, comprising PRMT1, PRMT2, PRMT3, PRMT4, PRMT6, and PRMT8, while type II PRMT5/7 symmetrically methylates arginine residues, and type III solely monomethylate the arginine residues. PRMT5, in association with its partner MEP50 or WDR7 shows optimum activity upon association, holds prominence as the primary member of type II PRMTs ^(10).^ On the other hand, EZH2 serves as the central subunit of polycomb repressive complexes. The H3K27me3 is considered a significant epigenetic process during cell fate determination and development, signify the EZH2, and show extensive associations with tumor initiation, progression, metastasis, metabolism, drug resistance, and immunity regulation ^(10, 14).^ Both EZH2 and PRMT5 have garnered considerable attentiveness as promising targets for novel drug development and are recognized as oncogenes. Recent studies have highlighted the functional collaboration of EZH2 and PRMT5, making them targets for combined therapy ^(8).^

The inhibition of EZH2 and PRMT5, is extensively reported using synthetic SAM-analogue compounds or their derivatives ^(12).^ In particular, selective inhibitors that compete with SAM binding site of EZH2 have been developed, this includes CPI-1205, CPI-0209, PF-06821497, Tazemetostat (EPZ-6438) and DS-3201, some of which are undergoing clinical trials ^(12–14).^ Similarly, PRMT5, targeted with inhibitors like sinefungin, GSK-3326595, and EPZ015666^(15–17).^ Ongoing clinical trials are evaluating the efficacy of these inhibitors against EZH2 and PRMT5, underscoring their importance as a crucial target for anticancer therapies. The development of inhibitors specifically tailored to PRMT5 and EZH2 is paramount, with natural compounds holding promise due to their potential for long-term use with minimal side effects.

EA, a naturally occurring polyphenol, is commonly present in various nuts and fruits, including berries such as raspberries, strawberries, and pomegranates ^(18, 19)^ and renowned for its robust antioxidant activity ^(20, 21)^. Additionally, EA has been extensively studied for its various health benefits, including its antiallergic, antiatherosclerosis, cardioprotective, hepatoprotective, nephroprotective, and neuroprotective properties ^(22).^ EA has demonstrated the ability to activate apoptosis, in cancer cells, effectively halting tumor growth ^(23).^ Additionally, it hinders tumor angiogenesis, the new blood vessels formation which is crucial for tumor metastasis and expansion ^(24, 25).^ Furthermore, it has been reported that it can inhibit or activate protein kinases ^(26)^ and interfere with NF-B activity thereby reduce proliferation of cancer cells^(27).^ Its anticancer efficacy enhances by its DNA damage shielding effect caused by carcinogens. Studies have shown that EA has anti-tumor properties by slowing down tumor cell growth, promoting tumor cell death, and limiting tumor cell spread and invasion through specific molecular pathways in different cancer types such as liver, breast, lung, and nasopharyngeal carcinoma ^(28).^ Collectively, EA by possessing its action in diverse signaling pathways and promised candidate for both cancer prevention and treatment ^(28–30).^**^)^** Highlighting the crucial role of EA in cancer chemoprevention, numerous studies have underscored its remarkable anticancer efficacy either alone or in combination with other compounds or drugs ^(31, 32).^ Despite of all outcomes, the precise binding target for EA has not been thoroughly investigated. The current study presents compelling evidence for the binding of EA to potential cancer targets, EZH2 and PRMT5 exhibiting high binding potential as demonstrated through molecular docking, Surface Plasmon resonance (SPR), and in vitro assays. The inhibitory potential of EA has been validated through both *in vitro* and *in vivo* studies targeting both EZH2 and PRMT5.

## 2. Materials and methods

### 2.1 materials

Ellagic-acid, S-adenosyl methionine (SAM), MTT: 3-(4,5-Dimethylthiazol-2-yl)-2,5Diphenyltetrazolium-Bromide, Dimethyl sulfoxide (DMSO), Recombinant proteins EZH2 (SRP0379) and PRMT5:MEP50 complex (SRP0145-25470 were purchased from Sigma-Aldrich United states. CM5 Sensor chip for SPR studies (cat no: 50-105-5511) was purchased from GE Healthcare Bio-Sciences (Cytiva life sciences-USA). The antibodies EZH2 antibody (#5246), Anti-PRMT5 antibody (#2252) H3 antibody (#4499), (H4) antibody (#2935), H3K27me3 (#9733), anti-beta actin antibody (#4967), Ki67 antibody (#D3B5) was purchased from Cell Signaling Technology (CST) USA. H4R3me2s Polyclonal antibody (A3718-050) procured from Epigentek NY-USA, FITC-Annexin V Dead cell/ Apoptosis kit (V13242) from Invitrogen-US. PRMT5 monoclonal antibody (MA-125470), EZH2 monoclonal antibody (MA5-15101) procured from ThermoFisher Scientific US. The antibodies for signaling studies BCL-2 (A0208), Bax (A0207), Bad (A1593), Beclin-1 (A7353), LC3 I/II (A19665), obtained from Abclonal Biotech Company US. Remaining all other chemicals are of higher standard unless otherwise mentioned.

### 2.2 Molecular docking studies

To perform molecular docking, we obtained protein structures of the Homo sapiens PRMT5:MEP50 complex with a SAM analog sinefungin,) and EZH2 from the RCSB Protein Data Bank (PDB IDs: PRMT5 - 4X60, 4X61, 4X63; EZH2 - 5HYN & 4Mi5). Subsequently, we prepared proteins by filling in missing residues using the Prime wizard in Schrödinger. The completed structures were then subjected to the Protein Preparation Wizard in Maestro to assign bond orders, hydrogens, and disulfide bonds. Additionally, all seleno-methionines were converted to methionine residues, and energy minimized using the OPLS3 force field. For docking simulations, we generated a receptor grid all directions of the active sites of the proteins, as determined by identifying interacting residues from the reference ligand bound to the crystal structure. Using the Glide program from Schrödinger Release 2019-3, we docked all prepared ligands alongside the reference molecule. Default parameters, including a Van der Waals radius of 1.0, were used for docking with extra precision (XP). The workflow included high-throughput virtual screening (HTVS), standard precision (SP), and finally, XP mode to ensure accuracy, optimal scoring, and visualization of the docked ligands. Subsequently, Glide results were analyzed to examine individual ligand poses, considering proximity (within 5 Å) of atoms, hydrogen bonds, and other contacts, as well as Glide XP scoring functions.

### 2.3 Protein-Ligand Kinetic Studies Using Surface Plasmon Resonance analysis

Surface plasmon resonance (SPR) analysis was conducted using the Biacore-T200 system (GE Healthcare) as described previously ^(33).^ Recombinant EZH2 and PRMT5-MEP50 complex (each 50µg) were diluted to 1mg/mL in 1x PBS pH 7.4 and immobilized onto a CM5 sensor chip using standard primary amine coupling. The amount of immobilized protein was estimated at 2500 RU per 2ng/mm². Reference surfaces (flow cells FC1 and FC3) were deactivated with 1M ethanolamine (pH 8.5), while activation of immobilization surfaces (FC2 for EZH2, FC4 for PRMT5-MEP50 complex) was achieved with a mixture of 200mM EDC and 50mM NHS. A stable baseline was established by flowing HBS-P buffer (10 mM HEPES pH 7.4, 150 mM NaCl, 0.005% P20+DMSO) over the immobilized proteins. Measurements were performed at 25°C with the same buffer flow. EA was prepared as a 10 mM working stock in PBS and injected over PRMT5-MEP50 complex and EZH2 at a flow rate of 30µl/min for 60 seconds at 25°C. Three-cycle kinetic analysis was conducted using triplicate injections. Sensograms were analyzed using BIA-T200 evaluation software version 2.1 (GE Healthcare). The relationship between protein concentration and Response unit (RU) value was calculated as 1000 RU = 1ng/mm² for surface concentration and 1000 RU = 10mg/ml for volume concentration. The dissociation rate constant (kd) was determined based on the Langmuir adsorption model.

### 2.4 Cell culture and *in-vitro* assays

For *in-vitro* assays, the human malignant cell lines MCF7 (breast), MDA-MB-231 (breast), PC-3 (prostate), Du-145 (prostate), HeLa (cervix), A549 (lung), Hep G2 (liver), and HEK293 (human embryonic kidney) were obtained from the National Centre for Cell Science (NCCS), Pune, Maharashtra, India. These cell lines were maintained at low passage numbers following standard sterile cell culture protocols. Cultures were grown in complete medium composed of DMEM (Dulbecco’s Modified Eagle’s Medium) high glucose (Himedia), supplemented with 10% (v/v) fetal bovine serum (Himedia) and 1x antibiotic-antimycotic solution (Himedia). Incubation was carried out at 37°C in a humidified atmosphere with 95% air and 5% CO2.

#### 2.4.1 Anti-proliferative Assay (MTT)

To study the anti-proliferative properties of EA, the viability assay was conducted as described previously ^(33, 34)^ 5000 cells were grown per well in triplicate on 96-well plates and incubated overnight. The media was then replaced with DMEM only. After incubation for 6 hours treated with various concentrations of EA, and incubated for 24 and 48 hours. Following the incubation period, the MTT assay was performed according to established protocols. Briefly, MTT solution was added and incubated for 3 hours, after which the MTT solution was aspirated, and 100µL of DMSO was added. Colorimetric quantification was then performed using a plate reader (Tecan). The percentage viability was calculated based on the optical density readings of the test and blank samples.

#### 2.4.2 Trypan blue dye exclusion assay

The trypan blue dye exclusion assay as previously described ^(33, 34)^ was employed in this study to assess cell viability following EA treatment. This assay is based on the principle that viable cells possess intact cell membranes that exclude the dye, while non-viable cells with compromised membranes take up the dye, rendering them visible under a light microscope. In detail, following EA treatment, a cell suspension was mixed with trypan blue dye. Subsequently, the mixture was examined under a light microscope to distinguish between cells that excluded the dye (viable cells) and those that took up the dye (non-viable cells). A graph was then plotted, representing the percentage of viable cells (with clear white cytoplasm) versus non-viable cells (with blue cytoplasm). To quantify cell viability, the following formula was applied:

% Cell Viability=Abs (Test sample)/Abs (control)×100%Cell Viability=Abs (control)Abs (Test sample) ×100. Additionally, the percentage of cell inhibition was calculated using the formula: % Cell Inhibition=100−percentage of Cell Viability %Cell Inhibition=100−% of Cell Viability This assay provides a straightforward and reliable means of assessing the impact of EA treatment on cell viability, offering insights into its potential inhibitory effects on cell proliferation or survival.

#### 2.4.3 EB/AO Double Staining and Fluorescent Microscopy for cell viability assessment

The EB/AO double staining assay, as previously outlined ^(33, 34)^ was employed in this study to evaluate cell viability. This assay utilizes two fluorescent dyes: Acridine orange (AO) and ethidium bromide (EB). AO is a cell-permeable dye that stains all cells, while EB can only penetrate cells with compromised membranes, such as necrotic cells or cells undergoing late-stage apoptosis. In this process, live cells with intact membranes will emit green fluorescence due to AO staining. In contrast, dead cells with compromised membranes will take up EB, resulting in red fluorescence. Necrotic cells, which have absorbed both dyes, will exhibit orange fluorescence, often resembling viable cells under microscopic observation, as there may be minimal changes in chromatin structure. MDA-MB-231 cells were treated with EA and then subjected to EB/AO staining after 24 hours of incubation. Observation was carried out using a fluorescent microscope to distinguish between live, apoptotic, and necrotic cells based on their fluorescence patterns. This technique offers precise insights into the impact of EA treatment on cell viability and the occurrence of apoptosis or necrosis within the cell population.

#### 2.4.4 Colony formation assay

The colony formation assay, based on established protocols ^(33, 34)^ with minor adjustments, was conducted to assess the impact of EA, on MDA-MB-231 cells. In this assay, 500 cells per well were seeded into 30mm culture dishes and allowed to adhere. Following attachment, cells were treated with varying concentrations of EA (0, 0.1, 1, and 10 µM) till 24 hours. After the treatment period, the media was replaced with serum-free media to inhibit further cell growth, allowing colonies to form over time. Once colonies became visible in the dishes, the media was carefully aspirated, and the dishes were washed with PBS to remove any non-adherent cells. Subsequently, 2-3 mL of a solution containing 6% glutaraldehyde and 0.5% crystal violet was added to each dish and incubated for 30 minutes. Following this incubation, the crystal violet-glutaraldehyde mixture was removed, and the dishes were rinsed with tap water by gently filling them. The dishes were then air-dried for 20 minutes. The experiment was performed in triplicate, and the number of colonies was counted. The results were plotted as the percentage of colony number relative to the control group. This assay provides valuable insights into the ability of EA to inhibit colony formation, reflecting its potential impact on cell proliferation and survival.

#### 2.4.5 Cell cycle and Apoptosis analysis through FACS studies

The 1.5 million cells per well were seeded in a six-well plate and treated with EA ranging from 0.1 to 10 µM, followed by a 24-hour incubation. Subsequently, the cells were treated with trypsin and stained with Annexin V-fluorescein isothiocyanate (FITC) with 100x dilution and propidium iodide (PI) at a concentration of 0.5 µg/ml for 15 minutes at room temperature. The FACScan (Becton Dickinson, San Jose, CA, USA) instrument was used to analyses the fluorescence intensities of Annexin V-FITC and PI The gating was carried as follows: Annexin V (+)/PI (-) cells were classified as apoptotic cells, Annexin V (+)/PI (+) cells had undergone secondary necrosis, and Annexin V (-)/PI (+) cells were considered necrotic cells. The obtained results were analyzed using FlowJo software.

To analyze the cell cycle, cells were trypsinized after treatment and fixed with 70% ethanol overnight at - 20°C followed, stained with RNase-A at a concentration of 100µg/ml and Propidium Iodide (PI) at a concentration of 50µg/ml for 15 minutes before flow cytometric analysis using FACScan (Becton Dickinson; San Jose, CA).

#### 2.4.6 Acidic Vesicular Organelle (AVO) staining by Acridine orange

Acidic Vesicular Organelle (AVO) formation identification by Acridine orange staining was performed as mentioned previously by ^(33)^ Acridine orange (AO) serves as an indicator for the formation of acidic vesicular organelles (AVOs), emitting green fluorescence throughout the cell except in acidic compartments, predominantly at late autophagosomes, where it emits red fluorescence. This characteristics of AVO development is observed during autophagy, indicating the maturation of autophagosomes. The only mature/late autophagosomes exhibit acidity. The intensity of the red fluorescence correlates with the number of AVOs present in autophagic cells. Following treatment with the indicated concentration of EA, the cell media was replaced with fresh media containing 5 µg/mL acridine orange and incubated for 10 minutes at 37°C before being observed under a fluorescent microscope.

### 2.5 Western Blotting

The cells were subjected to treatment with varying doses of EA (0-10 µM) for the indicated time. Later, the cells were washed with PBS and lysed in RIPA buffer supplemented with protease inhibitor cocktail, 1 mM PMSF and 1 mM DTT 4°C. The cell lysate was centrifugation at 10,000 rpm for 10 minutes at 4°C followed by protein estimation using Bradford’s protocol, BSA as a standard. The equal amounts of protein sample was loaded to SDS-PAGE (10%) as Laemmli method. After completion of electrophoresis protein bands were transferred to nitrocellulose membranes and blocked with 5% skimmed milk in Tris-buffered saline containing Tween-20. The respective primary antibodies are incubated at specified dilution as mentioned by manufacturers. The HRP-coupled secondary antibodies were used at a 1:10,000 dilution followed by development using ECL substrate from Bio-Rad Laboratories. The band image was captured using a Versadoc Imaging System and Image Lab 5.1 software (Bio-Rad). β-actin served as the loading control.

#### 2.5.1 Preparation of Lysate from Tumor Samples

Tumors from both control and treatment groups were dissected and weighed and washed with chilled 1x PBS and chopped into pieces. Subsequently, added ice-cold RIPA buffer with protease inhibitor (300μl), tissue was homogenized using an electric homogenizer with continuous agitation for 1 hour at 4°C. After homogenization, the lysate was centrifuged at 12,000 rpm for 20 minutes at 4°C. The supernatant was carefully collected in a fresh tube kept on ice, and the pellet was discarded. Protein concentration was determined using the BCA kit method, and the lysates were subjected to western blot analysis following the protocol described in section 2.5.

### 2.6 In-vitro Methylation Assay and ELISA

The in vitro methylation assay was conducted following the protocol described by ^(33, 34).^ Briefly, 1µg of the substrate (Histone H4/H3 acid, extracted, along with 0.1 mMol/µl of S-adenosylmethionine (SAM), was combined in a total volume of 30 µl PBS. For ELISA ^(33)^, the wells of a microplate were coated with histones (50µg/ml) in 0.2 M carbonate/bicarbonate buffer (pH 9.6) and covered with a lid, then incubated overnight at 4°C. The wells were washed twice with 200µl of 1x PBS. Subsequently, 100 µl of appropriately diluted samples of PRMT5-MEP50 with SAM, EZH2 & PRMT5-MEP50 with EA (pre-incubated with drug for half an hour at 30°C) were added, and the reaction was carried out at 37°C for one and a half hours. After the incubation period, the reaction mixture was removed, and the plate was washed twice with 200µl PBS. Blocking was performed by adding 200µl of blocking buffer (5% BSA/PBS) and incubating at 37°C for 2 hours. Following incubation, the blocking solution was removed, and the plate was washed three times with 200µl PBS for five minutes each. Then, 100µl of diluted (1:10,000) detection antibodies (H4R3me2s or H3K27me3) were added, incubated for 2 hours at room temperature, and washed four times with PBS. Subsequently, 100µl of secondary antibody conjugated with horseradish peroxidase (HRP) at a 1:5000 dilution in blocking buffer was added. The plate was incubated at room temperature for 1-2 hours, washed three times with 200µl PBS for five minutes each after the secondary antibody incubation, and then 100 µl of substrate reagent TMB was added to each well. The plate was covered and incubated for 15-30 minutes, and the reaction was stopped by adding 0.2M H2SO4. The absorbance was measured at 450nm.

### 2.7 Animal Studies

Swiss nude mice (female, 4 weeks old) were obtained from CSIR-Centre for Cellular and Molecular Biology Hyderabad India. Animals were maintained in University of Hyderabad animal house facility. All procedures were approved by the institutional ethics committee of University of Hyderabad and conducted in accordance with guidelines. The cages were maintained at 22±4°C with humidity of 50% to 60% in a 12-hour light/dark cycle. Laboratory diet and drinking water were provided ad libitum, and the mice were allowed to acclimatize for one week. After acclimatization, the mice were divided randomly into three groups with seven mice per group: ^(1)^ Control group-1, ^(2)^ Control group-2, and ^(3)^ Treatment group.

#### 2.7.1 Tumor Xenograft

The MDA-MB-231 cell lines were obtained from ATCC, used to get xenografts as previously instructed by ^(33).^ Mice were injected MDA-MB-231 (2x 106) cells subcutaneously on both flanks. Tumor volume was measured every three days with help of digital vernier calipers and following formula was used for calculation: Tumor volume = width2 x length/2. Once the tumors reached a volume of 100mm3, EA (100mg/kg) treatment was initiated for 21 days. Tumor samples were then harvested for weighing and further analysis.

#### 2.7.2 Immunohistochemical Assays

Tumor tissues obtained from both experimental and control mice were fixed overnight in 10% neutral buffered formalin, embedded in paraffin, and sectioned. Hematoxylin and eosin (H&E) staining was performed following the protocol described by ^(33)^ to visualize cellular and tissue structures in detail. For other histochemical analyses, tissues were deparaffinized and subjected to immunohistochemistry (IHC) using anti-Ki67 antibodies to assess proliferative activity. The stained tissue sections were visualized under a microscope after staining with DAB.

### 2.8 Statistical Analysis

Statistical significance was determined using analysis of variance (ANOVA), with P < 0.05 considered significant for both treated and untreated samples.

## 3 Results

### 3.1 EA Interacts with PRMT5 and EZH2

A comprehensive library of natural molecule from various plant species carried as reported Nalla et al 2024 ^(33).^ The molecular docking was carried to find the interaction between EA and EZH2 and PRMT5-MEP50 complex. The binding patterns of EA were assessed across three structural forms of PRMT5-MEP50 (4x60, 4X61, 4X63) which was reported with different ligands: SFG, SAM, and SAHA, respectively, in case of EZH2 (5HYN, 4mi5) with reported ligand H3K27 peptide and SET domain. As depicted in Figure -1 In the case of PRMT5, SFG interacted with the protein at Tyr324, Glu392, and Glu444 through hydrogen bonds, along with a π-cation interaction at the Lys393 position. The interaction of EA with the PRMT5:MEP50 complex involved a π-cation interaction at Lys393 and formed three hydrogen bonds with residues Tyr324, Glu392, and Glu444 in the PRMT5 pharmacophore. Remarkably, EZH2 (5HYN) exhibited strong binding with EA at -10.46 kcal/mol, forming a minimum of five hydrogen bonds with residues Trp624, Ser664, Ala622, Tyr728, and Asp732, whereas when docked with in place of 4mi5, EA interacted with Leu671, Phe670, Phe72, and Ser667 residues. The interactions of EA with PRMT5 (4X60, 4X61, 4X63) and EZH2 (5HYN, 4mi5) are summarized in (Table 1).

### 3.2 Binding study of EA with PRMT5:MEP50 and EZH2 by Surface Plasmon Resonance

The binding affinity of EA with EZH2 nd PRMT5 was performed using SPR. Recombinant EZH2 and PRMT5-MEP50 were immobilized on the sensor surface of a CM5 chip, and immobilization sensograms are presented Figure S2. Real-time bimolecular interaction analysis was employed to evaluate the interaction and affinity of EA with EZH2 and PRMT5-MEP50. Binding kinetics was analyzed by varying concentrations of the ligand over the immobilized proteins surfaces (Figures 2a,2c). The results demonstrate that EA displayed strong binding affinity in the micromolar concentration range (200-1.56 nM) with both PRMT5-MEP50 and EZH2. The equilibrium constant (KD), was calculated, which represents the strength of biomolecule interactions, the KD value is 3.28 E-06 M and 6.54E-05 for EZH2 and PRMT5-MEP50 respectively. The graph depicting response versus concentration exhibited linearity (Figures 2b,2d). Analysis of the association constant (ka) and dissociation constant (kd) revealed that EA exhibited a higher affinity for EZH2 compared to PRMT5, consistent with the findings from molecular docking studies.

**Figure-1:**
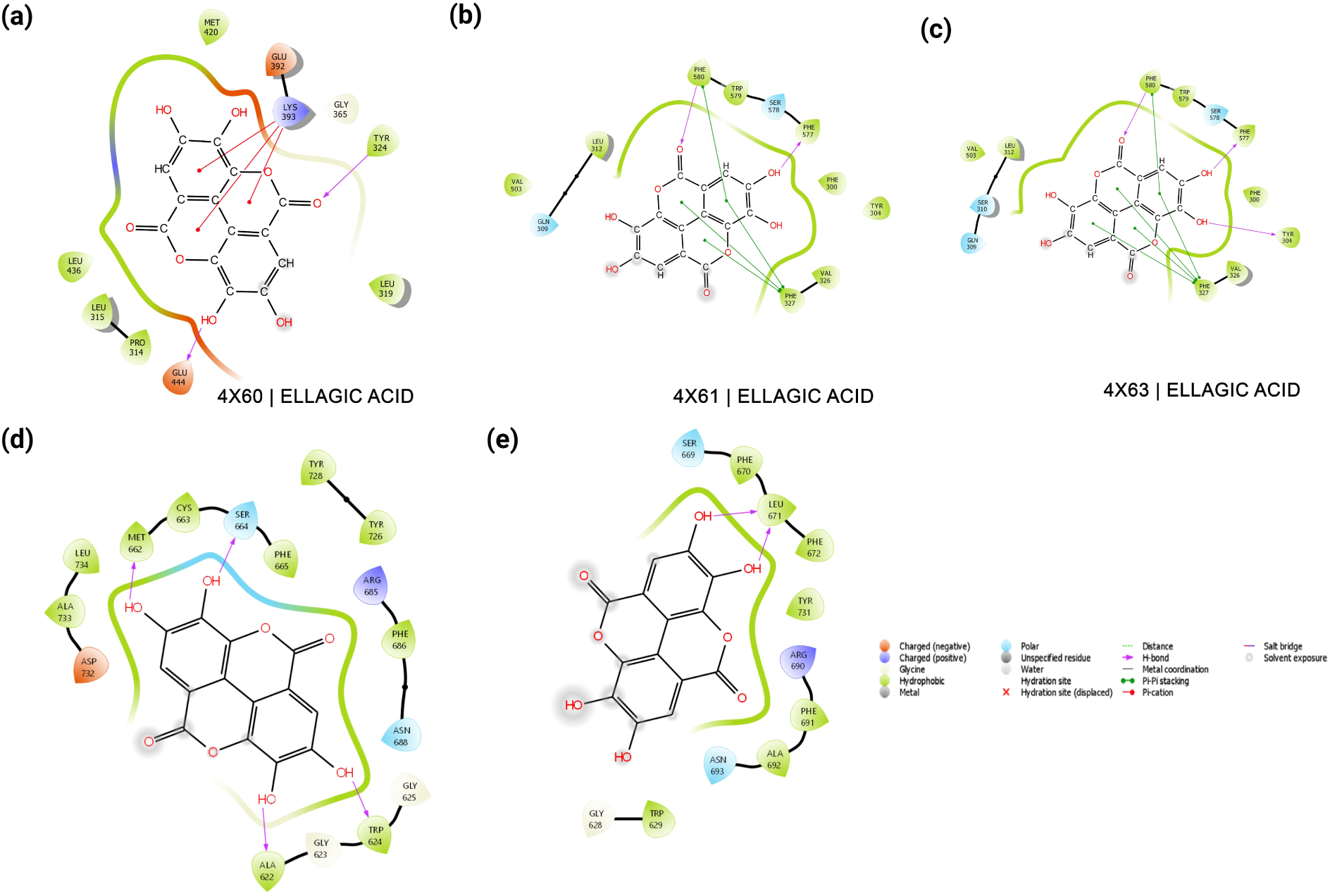
Interaction profiles of EA with the human EZH2 and PRMT5: MEP50 complex have been examined. (a-c) For the PRMT5: MEP50 complex (PDB IDs: 4X60, 4X61, 4X63), several key interaction features were identified. EA forms hydrogen bonds with specific amino acid residues, depicted by green dotted lines. These interactions, along with the ligand Sfg, are illustrated using a blue/black color ball and stick model. Non-bonded interactions are indicated by starbursts next to the residues. (d-e) For the EZH2 complex (PDB IDs: 5HYN and 4MI5), the ligands are centrally positioned. Hydrogen bonds are shown with purple arrows, with the arrowheads indicating donor-acceptor relationships. A significant π-cation interaction between EA, Sfg, and the lysine side chain is marked by a red arrow. Similar to the PRMT5 interactions, hydrogen bonds, amino acid residues, and the ligand Sfg are displayed using a blue/black color ball and stick model, while non-bonded interactions are represented by starbursts.

**Figure-2:**
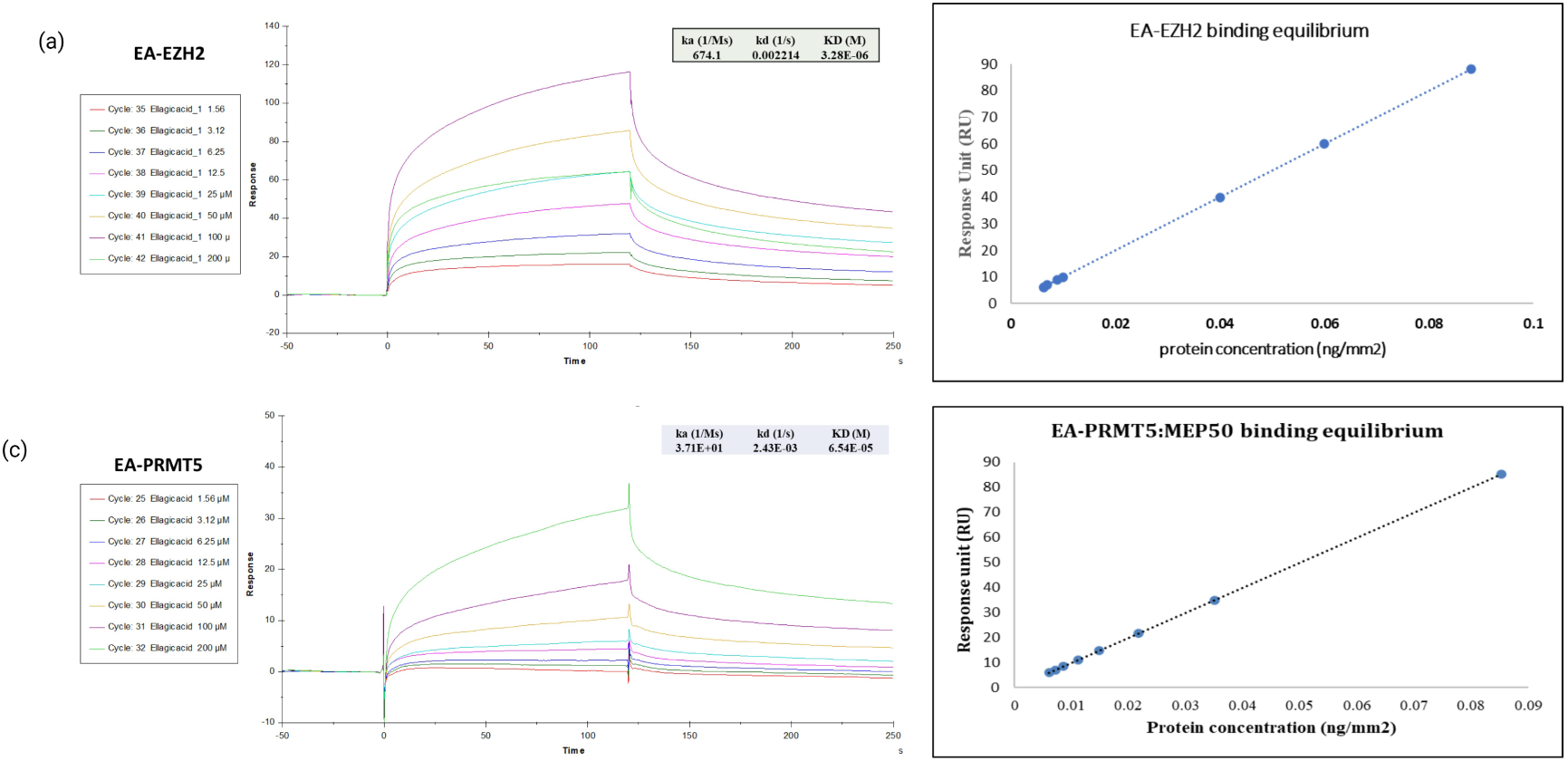
Surface Plasmon resonance analysis: Sensorgram analysis was performed to investigate the interaction of EA with EZH2 and the PRMT5:MEP50 complex using a CM5 sensor chip. (a) The dose-response sensorgram for EA interacting with immobilized EZH2 is shown. (b) The fitting of the response and concentration data was conducted using BIAcore T200 Evaluation software version 2.0. (c) The dose-response sensorgram for EA with immobilized PRMT5:MEP50 is illustrated. (d) The response and concentration data fitting for this interaction was also carried out using the same software. The plot of response units (RU) versus protein concentration demonstrated a linear relationship. The determined kinetic parameters are as follows: For EA with EZH2: association rate constant (ka) = 674.1, dissociation rate constant (kd) = 0.002214, equilibrium dissociation constant (KD) = 3.28E-06. For the PRMT5:MEP50 complex: association rate constant (ka) = 37.1, dissociation rate constant (kd) = 0.00243, equilibrium dissociation constant (KD) = 6.54E-05.

### 3.4 Inhibition of EA on growth of multiple cancer cell lines

Various human cancer cell lines, like MCF7 (breast), MDA-MB-231 (breast), DU-145 (prostate), PC-3 (prostate), A549 (lung), HeLa (Cervix), Hep G2 (liver), and HEK293 (human embryonic kidney), were employed in this study. The cells were treated with different concentrations of EA for 24 and 48 hours, and cell viability was assessed to determine the IC50 value (Figure S2). The trypan blue dye exclusion assay corroborated the results of the MTT assay, showing a decrease in viable cell numbers with increasing concentrations of EA (Figures 4b). Further, MDA-MB-231 cells were incubated with various concentrations of EA for 24 hours and examined for the levels of catalytic products of EHZ2 and PRMT5 EZH2, namely H3K27me3 and H4R3me2s (Figures 3a,3b). The results indicated a significant decrease in H3K27me3and H4R3me2s levels, significantly at concentration of 10 μM, compared to untreated cells. However, the protein levels of EZH2, PRMT5, H3and H4 largely remained unchanged. The β-actin was used as a loading control. In-vitro methylation assays of H3 and H4 followed by ELISA (Figure 3c) (1 µM, 10 µM) resulted in a dose-dependent decrease in H3K27me3 and H4R3me2s. Furthermore, AO-EtBr double staining revealed the presence of early-stage apoptotic cells, characterized by crescent-shaped or granular morphology, in MDA-MB 231 cells treated with EA after 24 hours of incubation, whereas no significant apoptosis was observed in the control group (Figure 4a). This observation was consistent with the results of the trypan blue dye exclusion assay and MTT assays (Figure 4b). In the colony formation assay, treatment with EA resulted in a dose-dependent decrease in colony formation compared to the control group (Figures 4c,4d). These *in-vitro* assays collectively demonstrated the potent inhibitory effect of EA on the activity of EZH2 and PRMT5:MEP50.

**Figure-3:**
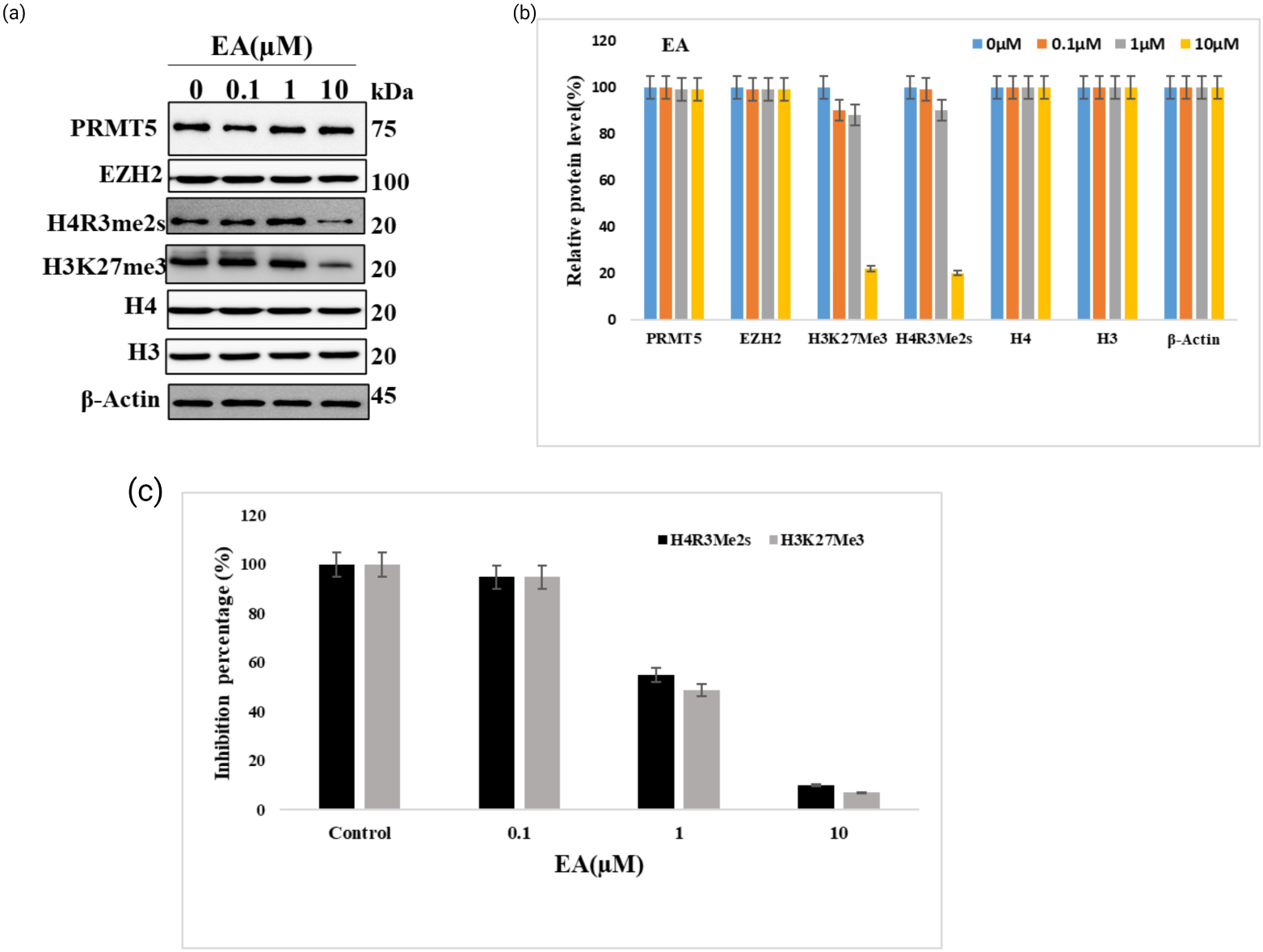
Impact of EA on H3K27me3 and H4R3mes methylation marks: MDA-MB231 cells were treated with increasing concentrations of EA (0.1, 1, 10 μM) for a duration of 24 hours. Following this exposure, protein lysates were prepared according to the described methods and subsequently analyzed for H4R3me2s and H3K27me3 levels using immunoblotting. β-actin was employed as a loading control. (a) Western blots. (b)Graphs depicting the intensity of the protein from western blot (c) *in-vitro* methylation was carried as described in methods and level of methylation was esteemed using ELISA. These assays utilized EZH2 and PRMT5-MEP50 enzyme complexes along with histones as substrates.

**Figure-4:**
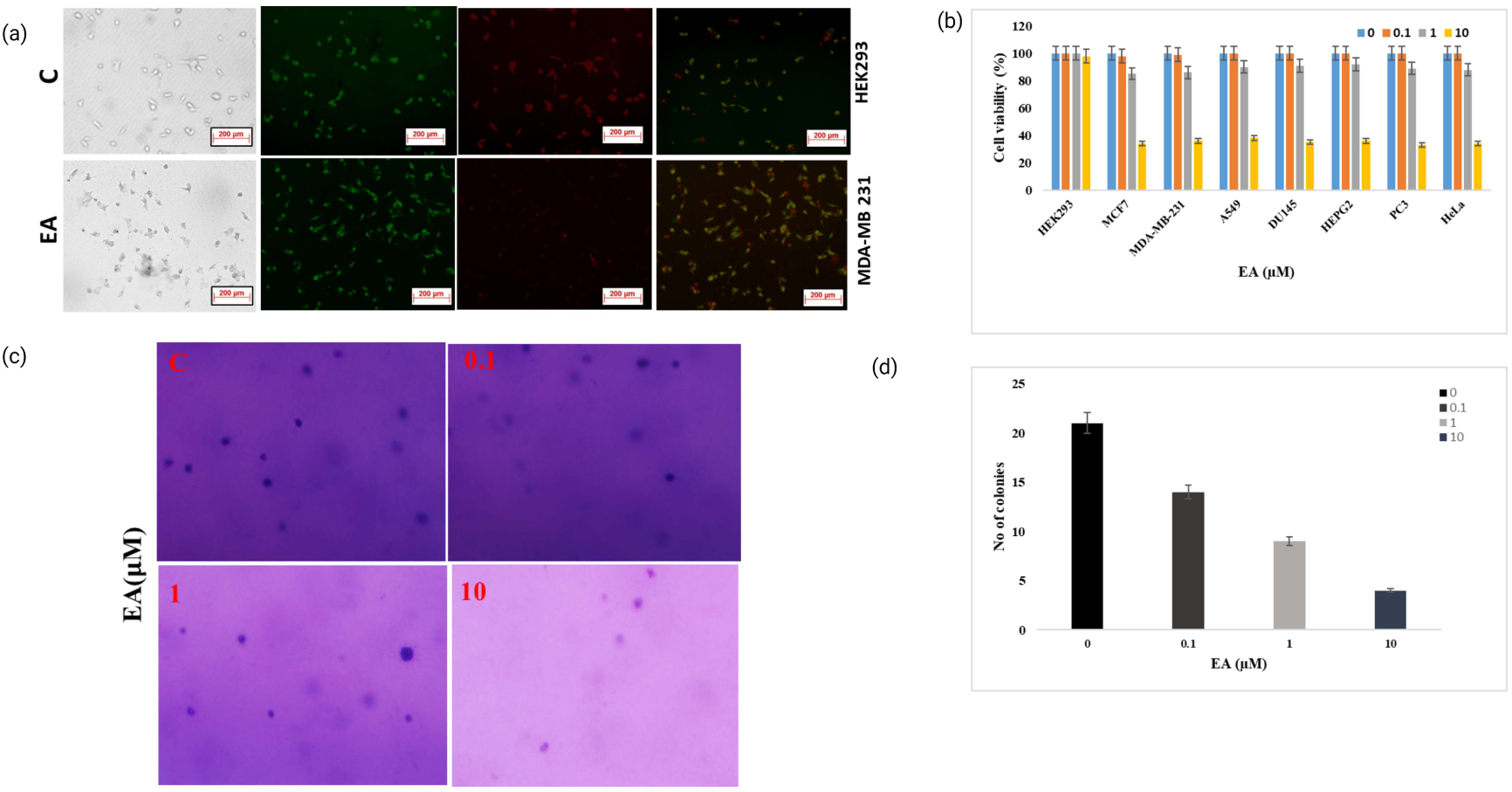
Effect of EA on Cellular Proliferation Dynamics: Cells were exposed to EA at specified concentrations and durations, followed by cell-based assays. (a) AO/EtBr assay: The cells underwent double staining with ethidium bromide and acridine orange (EB/AO), then were observed under a fluorescent microscope. Viable cells emitted green fluorescence (AO), while necrotic cells displayed predominant red fluorescence (EB). (b) Trypan blue dye exclusion assay: A549, MCF7, MDA-MB-231, Du-145, PC-3, HeLa, Hep G2, and HEK293 cells were treated with EA for 24 hours. Post-treatment, cells were incubated with Trypan Blue solution. Viable cells excluded the dye, whereas non-viable cells absorbed it, rendering them blue. Cell viability was evaluated under a microscope, and the outcomes were tabulated and graphed. (c & d) Colony formation assay: 500 cells were seeded and subjected to treatment per the experimental conditions. Following incubation, colonies were fixed and enumerated to evaluate clonogenic potential. These investigations offer valuable perceptions into the altered survival and proliferation of cells upon EA treatment.

### 3.5 Impact of EA on Apoptosis and Cell cycle

We investigated the effects of EA on apoptosis on MDA-MB-231 cells where cells were treated with varying concentrations (0, 0.1, 1, and 10 µM) of EA for 24 hours, followed by staining with Annexin V/PI to assess apoptosis levels via flow cytometry. Results revealed that a gradual increase in cells entering the late apoptosis stage with increasing concentrations of EA (Figures 5a,5b). Furthermore, we examined the potential impact of EA on cell cycle progression to assess cell cycle distribution and the distribution of cell cycle phases (Figures 5c,5d). Our findings indicated that both EA treatment resulted in G0/G1 arrest in a dose-dependent manner at concentrations of 1 µM and 10 µM compared to control.

**Figure-5:**
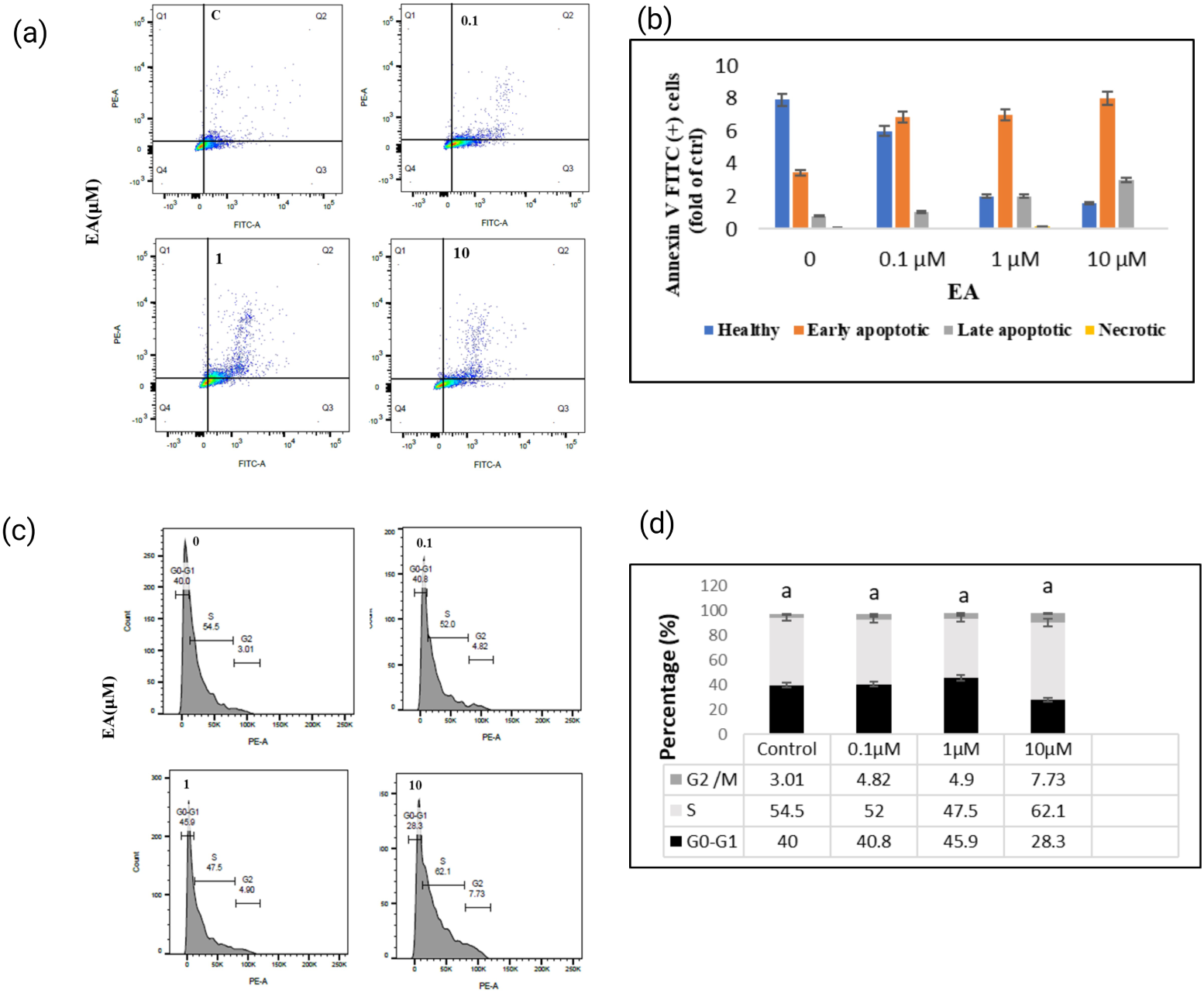
EA triggers apoptosis in breast cancer cells: MDA-MB231 cells were exposed to EA for a duration of 24 hours. Following treatment, cells were stained with Annexin V/propidium iodide (PI), and the proportion of apoptotic cells was assessed through flow cytometry. (a & b) Flow cytometry examination of MDA-MB231 cells treated with EA. (c & d) Analysis of cell cycle progression was performed on MDA-MB231 cells after 24 hours of treatment.

### 3.6 EA mediate the autophagy in cells

The AO-Etbr assay indicated the presence of apoptotic cells exhibiting characteristics such as crescent or granular shapes. To confirm whether the formation of granular cells was a result of apoptosis-mediated autophagy, we examined the formation of Acidic Vesicular Organelles (AVOs) followed by the EA treatment where the development of AVOs serves as an indicator for autophagy. We observed red fluorescent spots in cells treated cells, whereas control cells primarily displayed green cytoplasmic fluorescence (Fig 6a). Additionally, we investigated whether EA induced autophagy-mediated cell death in MDA-MB231 cells. To do so, we assessed the microtubule-associated protein light chain 3 (LC3),expression as a well-known marker for monitoring autophagy. Our findings demonstrated a dose-dependent transition of LC3 induced by EA (Fig 6b). Furthermore, to confirm autophagy induction, we evaluated the expression of Beclin-1 and BCL family proteins (BCL2, BAX, and BAD) (Fig 6b,6c).

**Figure-6:**
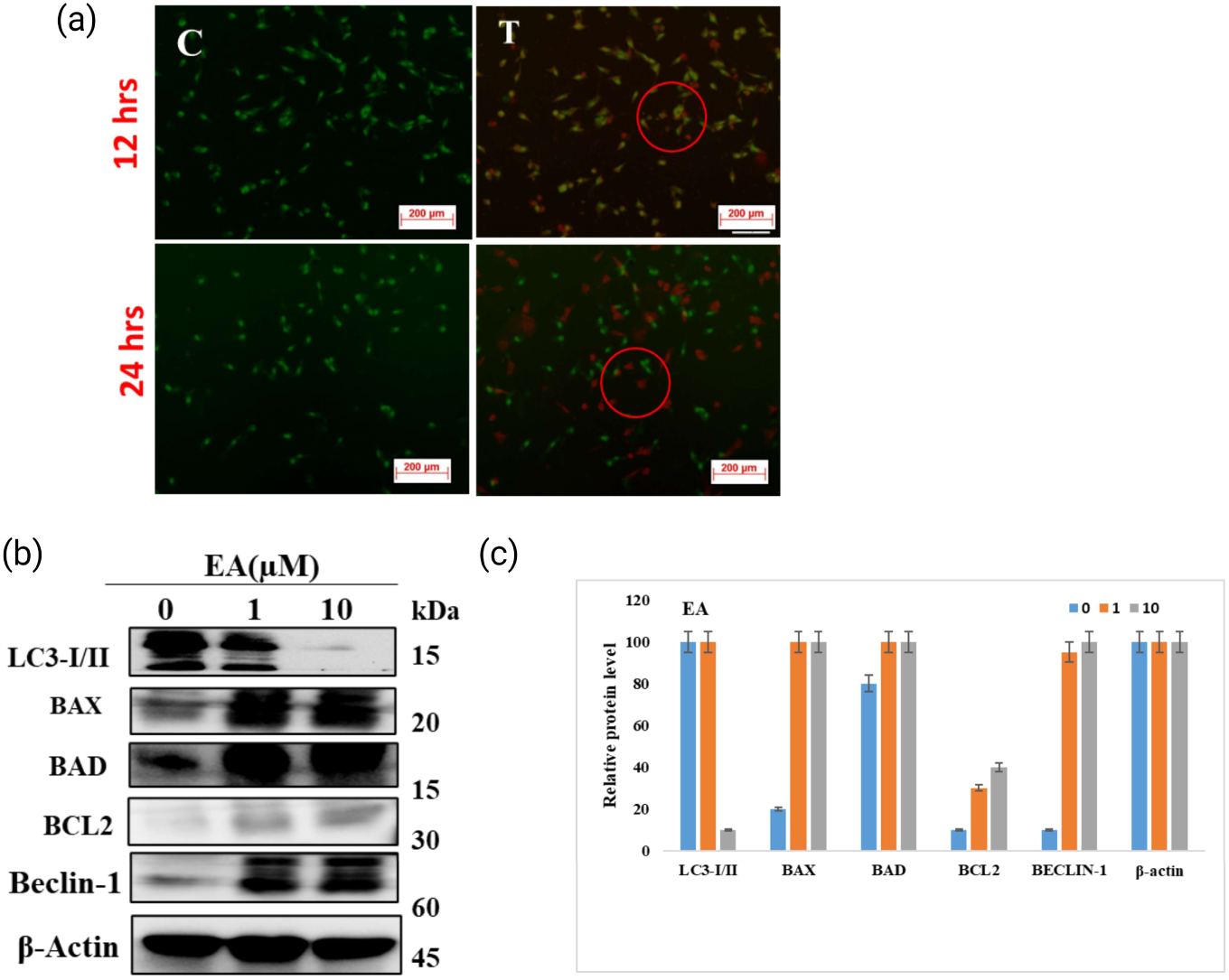
EA triggers autophagy in breast cancer cells: Following exposure of MDA-MB231 cells to EA for 24 hours, it was noted that EA induced autophagy in these cells. (a) Treatment with EA resulted in the formation of acidic vesicular organelles (AVOs) in cells. (b & c) Western blot analysis was performed to notice the expression of LC3 I/II and to evaluate the expression levels of Beclin-1, Bax, Bad and BCl2 proteins in MDA-MB231 cells treated upon with EA.

### 3.7 Impact of EA on tumor xenograft mouse model

To assess the anti-tumor effects of EA *in vivo*, we implanted subcutaneously MDA-MB231 cells into Swiss nude mice to generate tumor xenografts as explained in methods. After tumor development, we measured the tumor weight, followed with administration of EA at a dosage of 100mg/kg upto 21 days. The results indicated that a noteworthy reduction in tumor volume when compared to the control groups (Fig 7a-7c). Meanwhile, the body weight of mice in the control group showed a steady increase throughout the experiment period (Fig 7g). Furthermore, to evaluate the potential toxicity of EA, we performed H&E staining on tumor tissues. A comparison between control tumors and those treated with EA revealed notable differences. Control group tumors exhibited highly aggressive sarcomatous neoplastic cells with a vacuolated appearance and increased mitotic activity (Fig 8ai, 8aii). In contrast, treated tumors showed sections with only subcutaneous and muscular layers, lacking dermal and epidermal tumor regions (Fig 8aiii-8aiv). Moreover, the proliferation of cells was assessed using the Ki-67, a proliferation marker. IHC analysis demonstrated that both EA significantly reduced Ki-67 levels in the treated groups compared to the control group (Fig 8bv-8bviii). Further to check the ability of EA on EZH2 and PRMT5 activity, we analysed the level of catalytic products of bothEZH2 and PRMT5, using western blot of tumors from control and EA treated groups. The results revealed a significant reduction in the H4R3me2s and H3K27me3 in tumors treated with EA when compared to control (Fig 7d,7f).

**Figure -8:**
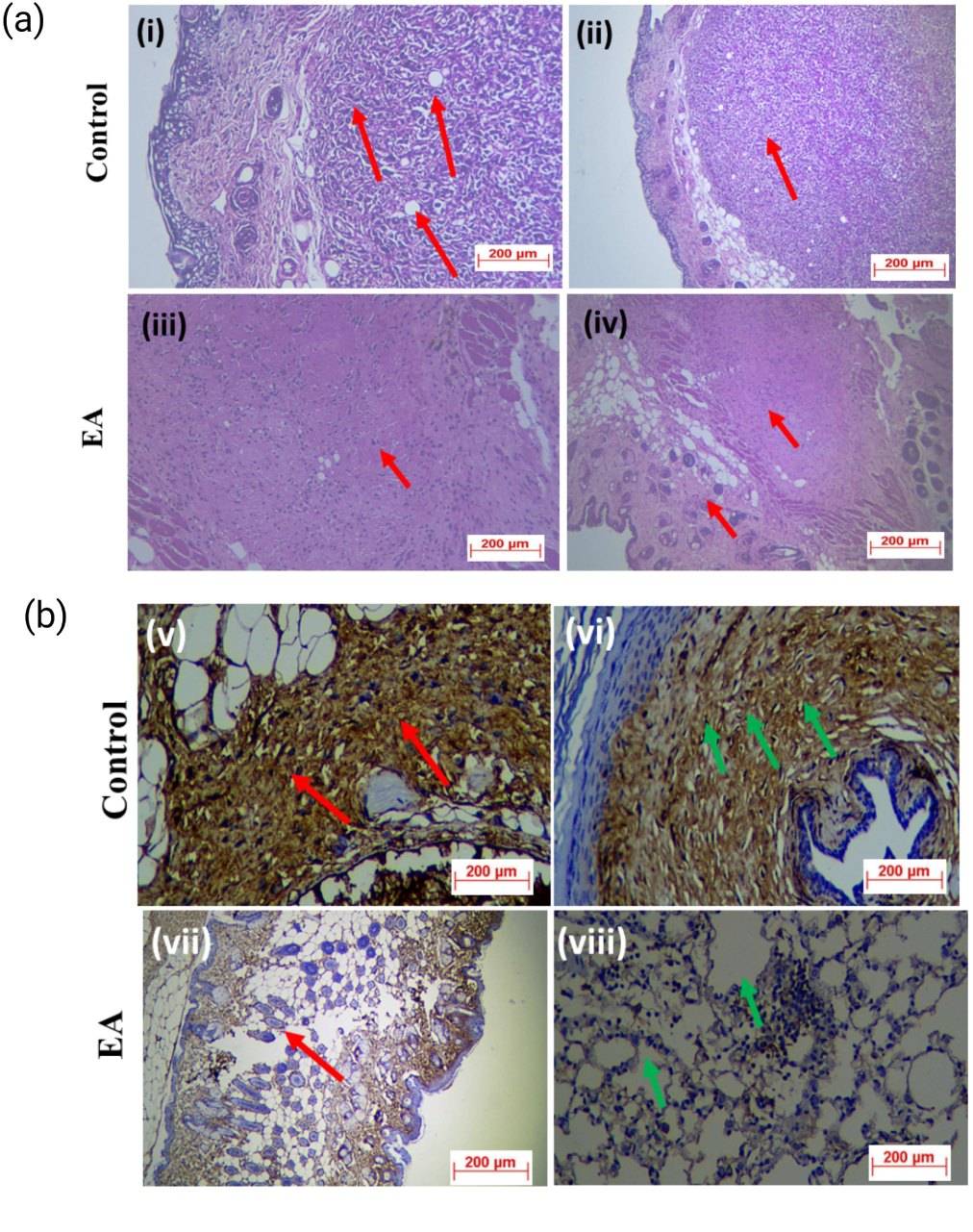
Immunohistochemistry (IHC) studies: post drug administration for 21 days, the mice were euthanized as per IEC rules, and tumors were collected from control and treated groups. Tumor samples underwent histopathological examination by an expert for IHC studies. A comparison was drawn between control tumors and those treated with EA. In the H&E staining analysis (8a (i, ii)), control tumor samples exhibited aggressive neoplastic sarcomatous cells forming nodules in the subcutaneous region, invading into the dermal region, as evidenced by mitotic figures (red arrows). Conversely, EA-treated tumor samples (8a(iii-vi)) displayed only subcutaneous and muscular layers, devoid of dermal or epidermal tumor regions. The subcutaneous region appeared normal, with no metastatic invasion of neoplastic cells (red arrow), and the muscular region showed no metastatic invasion (green arrow). Regarding Ki67 expression (8b (v-viii)), control tumor samples demonstrated intense expression of Ki67 in the subcutaneous tumor mass (red arrows), with neoplastic cells invading into dermal and dermal hair follicles (green arrow) (8b viii). In contrast, tumor samples treated with EA showed 10-20% expression of Ki67 in pleomorphic anaplastic epithelial cells in the epidermal layer (red arrows). Overexpression of Ki67 was also noted in stromal tissue in the dermal and subcutaneous regions (green arrows).

**Figure-7:**
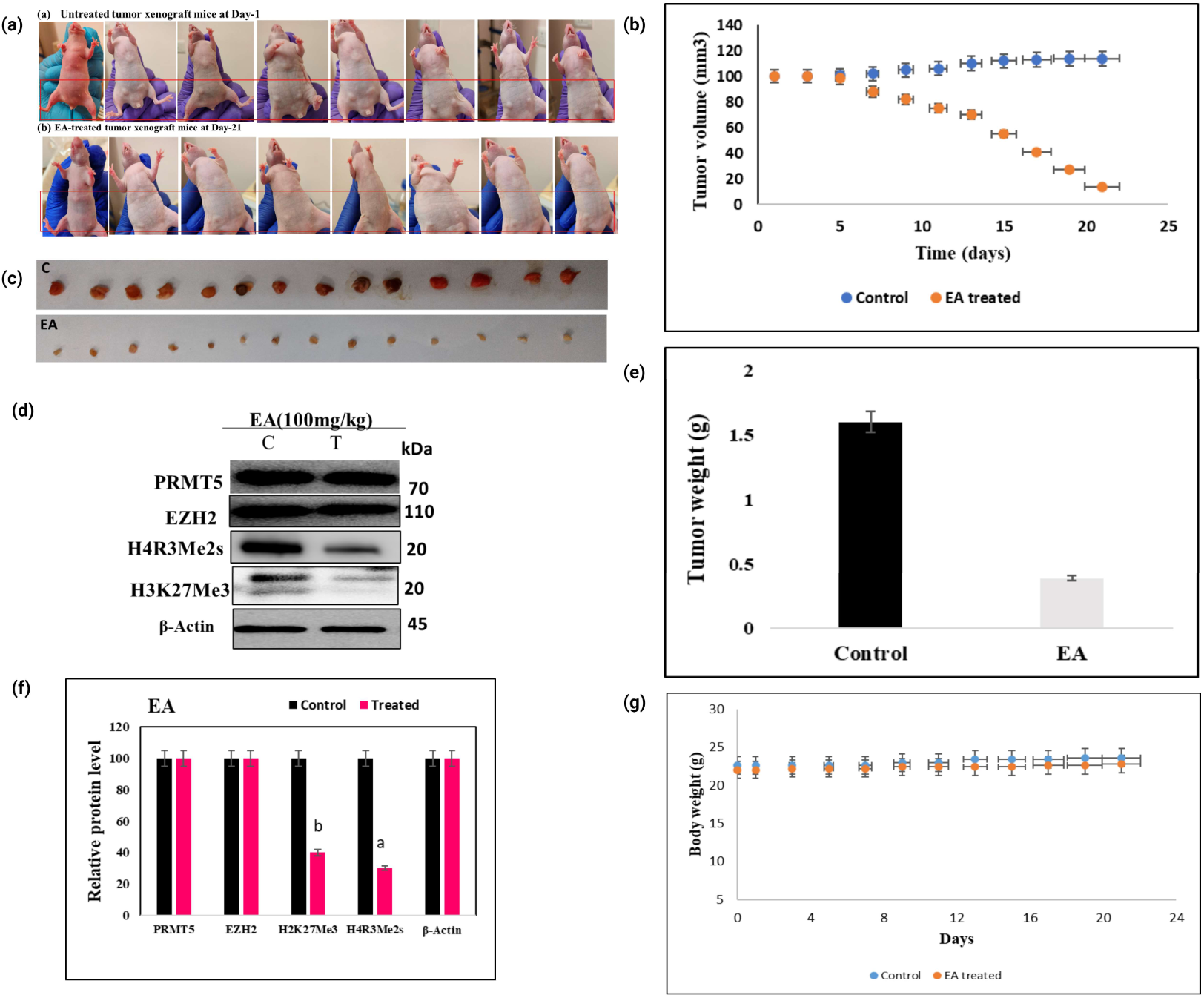
In-vivo xenograft model studies: Mice were randomly assigned to one of three groups, each containing 7 mice: ^(1)^ control group-1, ^(2)^ control group-2, and ^(3)^ treatment group. Subcutaneous injections of MDA-MB231 cells were administered to both flanks of the mice. Tumor volume was monitored every three days, and once reaching 100 mm3, oral administration of EA (100mg/kg) commenced for the treatment group, continuing for 21 days. (a) Comparative analysis of mice treated with control mice. The initial mouse represents the control, while subsequent mice (2nd to 7th) developed tumors on the first day of treatment. (b) Graphical representation comparing tumor volumes over the 21-day period between control and treated mice. (c) Evaluation of tumor growth, with the upper panel presenting data from control animals and the lower panel showing data from the treatment group. (d) Tumor lysates western blot analysis from the treatment group, focusing on levels of EZH2, PRMT5, EZH2, H3K27me3, H4R3me2s, and β-actin as a loading control. (e) Assessment graph illustrating tumor weights between control and treated groups. (f) Comparison of protein intensity levels observed in western blot studies between control and treated groups. (g) Graphical representation of body weight measurements of control and treated mice.

## Discussion

The epigenetic, processes like DNA methylation and histone modifications such as methylation, phosphorylation, acetylation, sumoylation and ubiquitination paved the major contribution towards the formation of cancer ^(35)^. These alterations impact chromatin structure and various pathways associated with the development of cancer. In recent decades, tremendous progress has been made in developing therapeutic strategies for cancer, with several clinically approved drugs targeting key epigenetic regulators such as DNA methyltransferases (DNMTs), histone deacetylases (HDACs), histone methyltransferases (HMTs), histone demethylases (HDMs), and bromodomain and extra-terminal proteins (BETs) ^(36–42).^ Notably, inhibitors of DNMTs (DNMTi), HDACs (HDACi), and BET proteins (BETi) have received approval for specific cancers, reflecting their efficacy in clinical settings ^(15).^ Furthermore, there is growing interest in developing inhibitors targeting PRMT5 and EZH2 due to their versatile roles in various cancers ^(16, 17, 43–46).^ Both EZH2 and PRMT5 overexpressed in various cancers including breast, prostate, endometrial, liver, small cell lung cancer ovarian, melanoma, glioblastoma, pediatric glioma, bladder, and lymphomas ^(16, 43–48)^ hence PRMT5 and EZH2 are potent targets for cancer therapeutics. ^(7, 8).^ Histone modifications regulate chromatin accessibility and modulate gene expression patterns. Plethora of synthetic small molecule as inhibitors of EZH2 and PRMT5 are different stages of development. ^(47, 77).^ Despite of these the advancements in therapeutic strategies and treatments lead to severe side effects in cancer patients. As a result, there is a global effort among researchers to explore the anti-cancer effects of natural compounds with minimal adverse reactions. A wide range of phytocompounds, encompassing curcumin, resveratrol, brazilin, catechin, quercetin, EGCG, and EA, have emerged as potential agents to target epigenetic regulatory proteins ^(42).^ These natural compounds have demonstrated the ability to interact with various key epigenetic regulators, including DNMT1 and HDAC1, within their catalytic pockets ^(48).^ These interactions underscore the potential of phytocompounds to directly influence the activity of crucial epigenetic enzymes involved in cancer progression ^(49, 50).^

In present study we demonstrated binding potential of EA to the catalytic pocket of EZH2 preferentially than the PRMT5. It has been reported that functional properties of EA towards the various action towards like antioxidant enzymes ^(8, 9).^ like catalase (CAT), superoxide dismutase (SOD), glutathione peroxidase (GPx), and glutathione-S-transferase (GST) ^(51, 52).^ In another study demonstrated suppression of cyclooxygenase (COX-2) and nuclear factor-kappa B (NF-кB) ^(53).^ Additionally, it modulates various signalling pathways, like phosphoinositide 3-kinase, nuclear erythroid 2-related factor 2 (Nrf2) and glycogen synthase kinase 3 beta (GSK-3β) in endometrial cancer. (^23, 66, 67, 76^).

Previous *in vitro* studies have highlighted the beneficial effects of EA against colorectal, breast, prostate cancers, leukaemia, lymphoma, and melanoma ^(53–58).^ However, the molecular targets of EA have not been explored, present study molecular docking experiments confirmed interaction of EA with EZH2 more strongly than PRMT5 with binding energy -10.46kcal/mol. As depicted in Figure 1, EA exhibited similar interaction patterns as observed with known ligands such as 5HYN and SFG of EZH2 and PRMT5 respectively. Notably, EZH2 showed strong interaction with EA, with five hydrogen bonds withTrp624, Ser664, Ala622, Tyr728, and Asp732. In case of PRMT5, EA formed hydrogen bonds at Tyr324, Glu392, and Glu444 residues, in addition π-cation interaction at the Lys393 position as shown by ligand SFG ^(82, 83).^ This observation is supported by SPR binding studies, the obtained association rate constant (ka) equilibrium dissociation constant (KD) and dissociation rate constant (kd), values are as follows: For EA with EZH2: Ka = 674.1, kd = 0.002214, KD = 3.28E-06. For EA with PRMT5 complex: Ka = 3.71E+01, kd = 2.43E-03, KD = 6.54E-05. Interestingly, when compared to previous reports, EA showed potent binding interactions with EZH2 than the well-studied synthetic molecules like CPI-1205, CPI-0209, Tazemetostat (EPZ-6438), PF-06821497, and DS-3201, some of which are under clinical trials ^(64–66).^ In vitro methylation assays, followed by ELISA has confirmed the reduced levels of specific histone *in vitro* (H3K27Me3 and H4R3mes) as well as in MCF7 and MDMB231 cell after 24hr treatment further validated the inhibitory protentional of EA. ^(21, 32, 49).^

Polyphenolic compounds exert their anti-cancer properties through various mechanisms, including antioxidant, anti-inflammatory, and anti-proliferative actions ^(80, 81).^ Additionally, it can influence cellular signalling pathways, induce cell-cycle arrest, and promote apoptosis ^(49).^ Numerous anticancer molecules have been found to induce autophagy ^(59).^ Studies have reported the inhibition or down regulation of both EZH2 and PRMT5, induce the apoptosis, autophagy, and cell cycle arrest ^(60–62).^ Autophagy exhibits a dual role in cancer, capable of both supporting cancer cell survival and inducing cell death ^(79).^ The inhibition of EZH2 by 3-deazaneplanocin A (DZNep), a specific EZH2 inhibitor, induce apoptosis in NRK-52E cells through the H3K27me3 in its promoter ^(63).^ Similarly, the dereased activity of EZH2 by EA may lead to autophagy and apoptosis. In another study in colorectal cancer cells, EZH2 down regulation induced autophagy and apoptosis, while siRNA mediated inhibition of EZH2 decrease the proliferation and activate the apoptosis by inducing G0/G1 cell cycle arrest in SW620 cells ^(64).^ In human lung cancer cells, inhibition of PRMT5 with GSK591 or shRNAs induces apoptosis and autophagy, accompanied by decreased Akt/GSK3β phosphorylation and reduced level of cyclin D1 and E1 ^(66).^ The relation of PRMT5 activity and cancer has been reported by many researchers. For instance, the PRMT5 knockdown enhances cell pyroptosis in multiple myeloma ^(67),^ and modulate the expression of cell cycle and apoptosis related genes by modifying p53, KLF4and E2F-1 ^(15^). In another study it has been showed that PRMT5 inhibition leads to increased apoptosis and decreased cell growth ^(68).^ PRMT5 activity influences different stages of autophagy, controlling autophagy and tumorigenesis through specific target modulation ^(69).^ Brobbey et al (2023) reported in triple-negative breast cancer cells that methylation of ULK1 catalyzed by PRMT5 at R532, suppressed the ULK1 activation and affected autophagy ^(70).^ A well-known epigenetic inhibitors Trichostatin A suppressed cervical cancer cell proliferation and induced apoptosis and autophagy through regulation of the PRMT5/STC1/TRPV6/JNK axis ^(71).^ The global reduction in the H4R3me2s and H3K27me3 repressive histone methylation marks may induce apoptosis and autophagy, as observed in the present study. These marks are associated with cell cycle arrest (71, 78). This modification directly or indirectly involved in the regulation of different cell cycle regulators like cyclin D1, cdk4, cdk6, p21, and p27 in pancreatic cancer cells ^(72),^ and EGFR/cyclinD1 signaling in lung cancer ^(73),^ and WAF1/p21 in prostate cancer ^(60, 69–71).^ These evidences substantiate our results that **EA** mediated reduced level of above histone modifications may be contributes to cell cycle arrest, (G0/G1, G2/S, G2/M) in MDA MB231 cells.

It has been reported in *vivo* model that role of EZH2 and PRMT5 in synthetic small molecules reduced the tumor size as well as recover from the cancers (10, 11, 14) In the present study upon treatment with EA in MDA-MB231-induced xenografts mouse model significantly reduce the tumor size. The reduction in tumor size is in concurrent with reduced level of H3K27me3 and H4R3mes however no rection in EZH2 and PRMT5 level observed in western blots of tumor samples. Perhaps we do not have direct evidence to link the reduced level of tumor size and with decreased level of these histone marks through any signaling pathway or promoter occupancy of these histone marks. This reduction was supported by a substantial decrease in the proliferative marker Ki76, although a direct link between Ki67 and reduced levels of H4R3mes and H3K27me3 was not evident. The inhibitory effects of EA, of oncogenic PRMT5 and EZH2 suggest its potential as a potent anti-cancer agent. Finally, we demonstrated inhibitory potential of natural compound EA towards the EZH2 and PRMT5 using *in silico*, *in vitro* and *in vivo* model. These findings represent future therapeutic potential of natural molecules as an epidrug.

**Table1:**
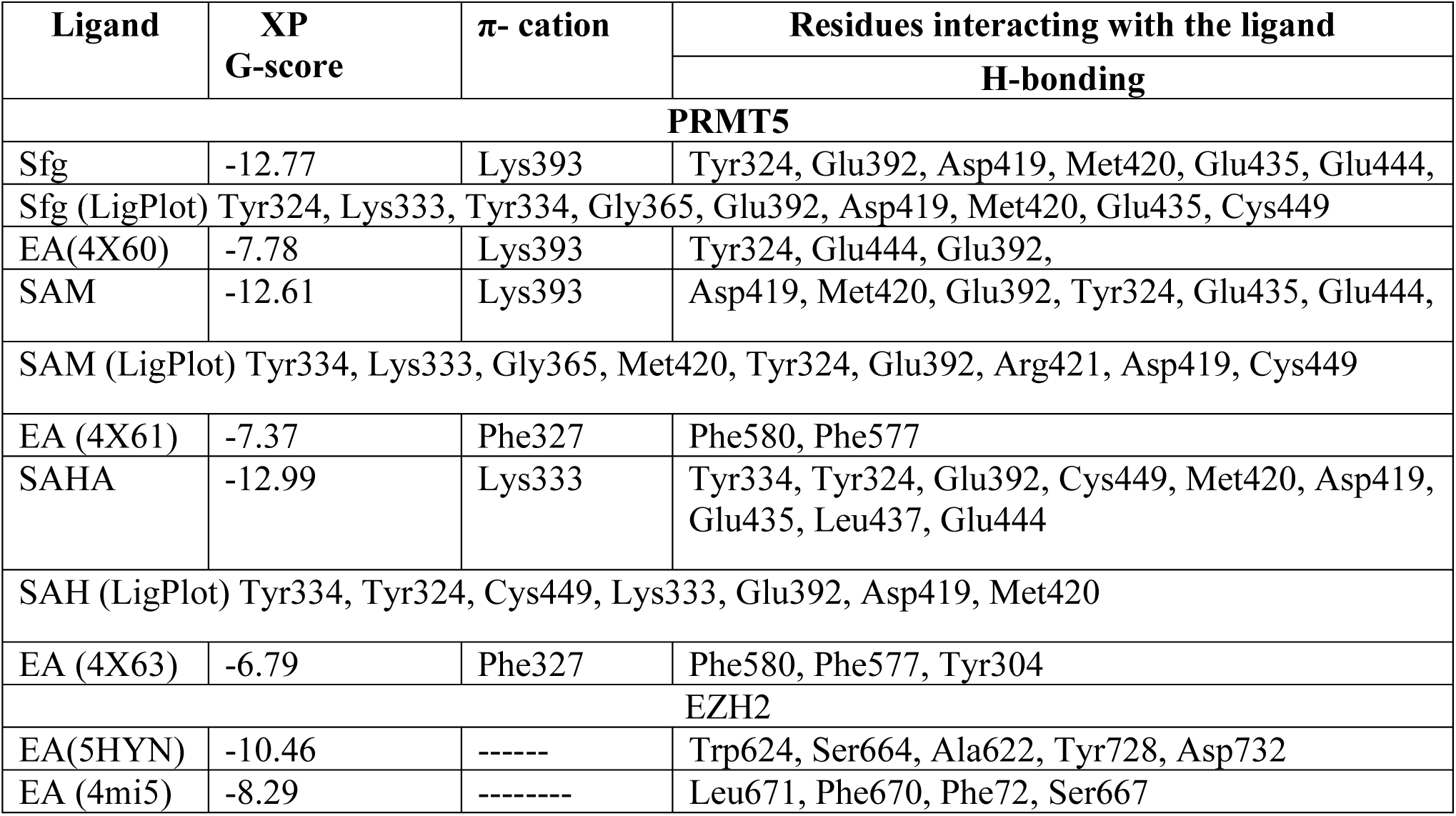
Interaction profiles of EA with human EZH2 and PRMT5: MEP50: were analyzed using several protein complexes, specifically 5HYN and 4MI5 for EZH2 and 4X60, 4X61, 4X63 for PRMT5: Molecular docking studies were conducted with these complexes. Additionally, the GlideXP docking scores of EA with Sfg, SAM, and SAH were evaluated and are included.

## Author contributions

NKK: Performed the experiments and wrote the manuscript; BC: Performed the initial screening of compounds JP; Helped and performed molecular docking, KG: Conceptualized the *in-silico* work AS: Helped in *in-silico* and molecular docking studies AP; Provided molecular docking platform; CJ; Provided the molecular docking software platform and critical comments, BM: Helped in animal studies, and given critical comments. SRK: Conceptualized, designed the experiments, interpreted the data, supervised the entire work, and wrote the manuscript.

## Acknowledgement

Authors are thankful to University of Hyderabad Grant (UoH-IoE-RC3-21-008) for financial support. Authors also acknowledge the DBT Builder and DST-FIST/UGC-SAP departmental facilities. KKN is thankful to Department of Science and Technology, Govt. of India (DST/INSPIRE/03/2017/000035 & IVR No-201700011820) for DST-INSPIRE Fellowship and AS thankful to DST-INSPIRE fellowship ((DST/INSPIRE/03/2018/000039 & IVR No-201800024300), provided by Department of Science and Technology, Govt. of India Heartfelt thanks to Ms Mounika and Dr K Praveen (Cytiva) for helping in SPR experiments.

## Abbreviations

ADMA: Asymmetric dimethylarginine
AVO: Acidic vesicular organelles
(AO): Acridine orange
BAD: BCL2 associated against of cell death
BCL-2: B-cell lymphoma-2
Bax: Bcl2 associated protein X
BSA: Bovine serum albumin
DMSO: Dimethyl sulfoxide
DMEM: Dulbecco’s Modified Eagle’s Medium
DNMT: 
EB: Ethidium bromide
EA: Ellagic acid
EDC: N-ethyl-N’-dimethylaminopropyl carbodiimide
EGCG: Epigallocatechin-3-gallate
EZH2: Enhancer of Zeste homolog 2
ELISA: Enzyme linked immunosorbent assay
FC: Flow cell
FBS: fetal bovine serum
H3K27: H3 residue on lys27
H3K27me3: Trimethylated lysine 27 on histone H3
H4R3: H4 residue on Arg3
H4R3me2s: Symmetrically dimethylated arginine 4 on histone H4
HMTs: Histone methyltransferases
HTVS: High-throughput virtual screening
HRP: Horseradish peroxidase
H& E: Hematoxylin and eosin
IHC: Immunohistochemistry
HDACs: Histone deacetylases
HDM: Histone demethylases
LC3: I/II Microtubule-associated proteins 1A/1B light chain 3B
MTT: 3-(4,5-Dimethylthiazol-2-yl)-2,5Diphenyltetrazolium-Bromide
MEP50: Methylosome protein 50
MMA: Monomethylarginine
NHS: N-hydroxysuccinimide
PRMT5: Protein arginine methyltransferase 5
PRMTs: Protein arginine methyltransferases
PRC2: Polycomb Repressive Complex 2
PBS: Phosphate buffer saline
RU: Response unit
SAM: S-adenosyl methionine
SDMA: Symmetric dimethylarginine
SPR: Surface plasmon resonance
WHO: World health organization

## Supplementary figures

**Figure S1:**
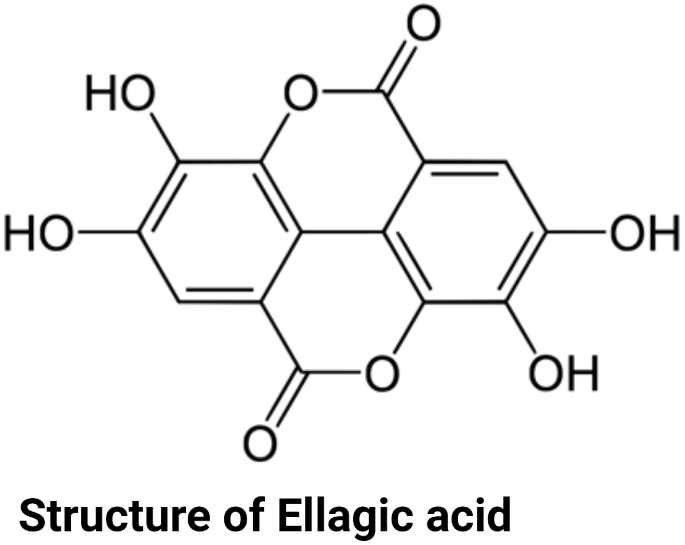
Chemical structure of EA (Mol wt. 302.19 g/mol, chemical formula: C14H6O8

**Figure S2:**
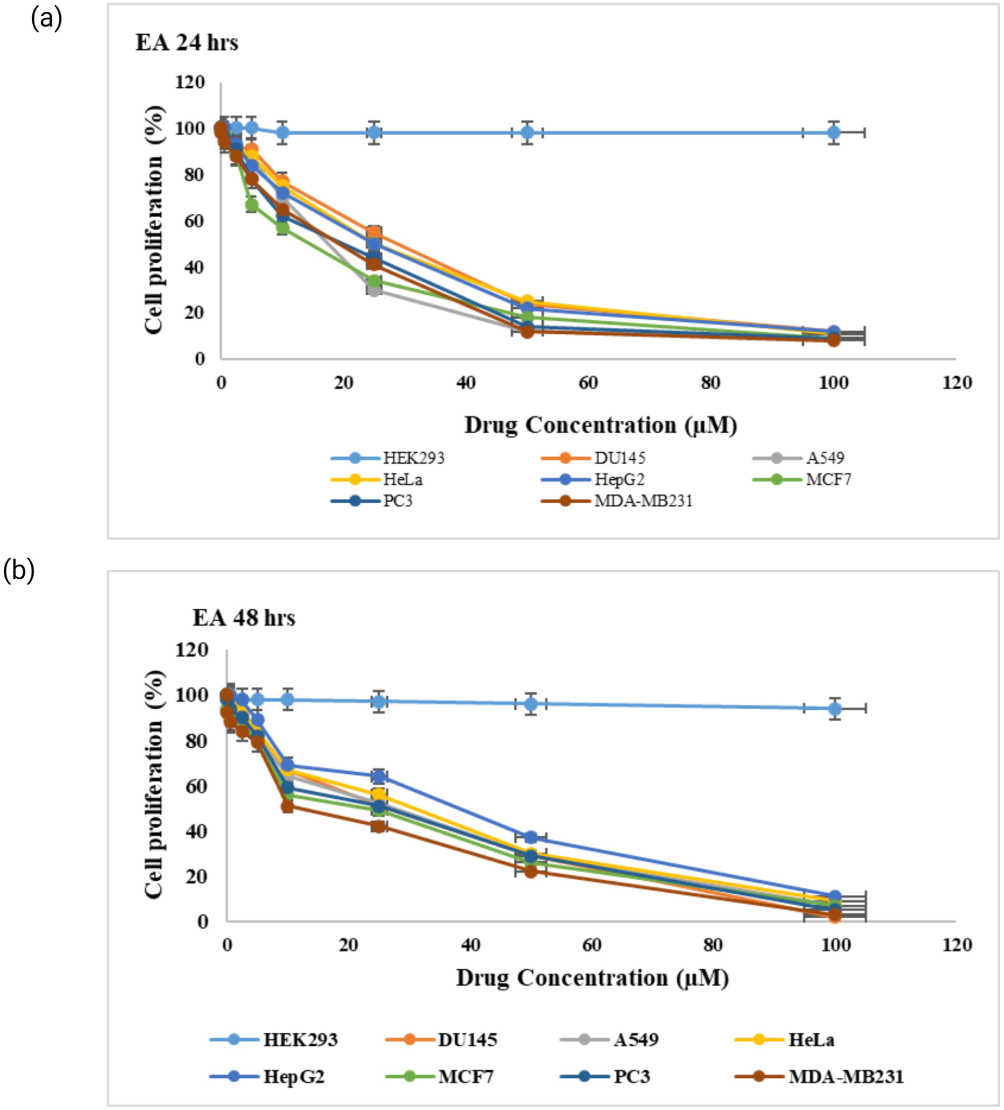
Impact of EA on human cancer cell lines: The human cancer cell lines MCF7 (breast), MDA-MB-231 (breast) Du-145 (prostate), A549 (lung), Hep G2 (liver), HeLa (Cervix), PC-3 (prostate) and HEK293 (human embryonic kidney) were exposed to different concentrations of EA for 24 & 48 hours for cyto-toxicity and determined Ic50 values and tabulated. (a) EA for 24 hours (b) EA for 48 hours

**Figure S3:**
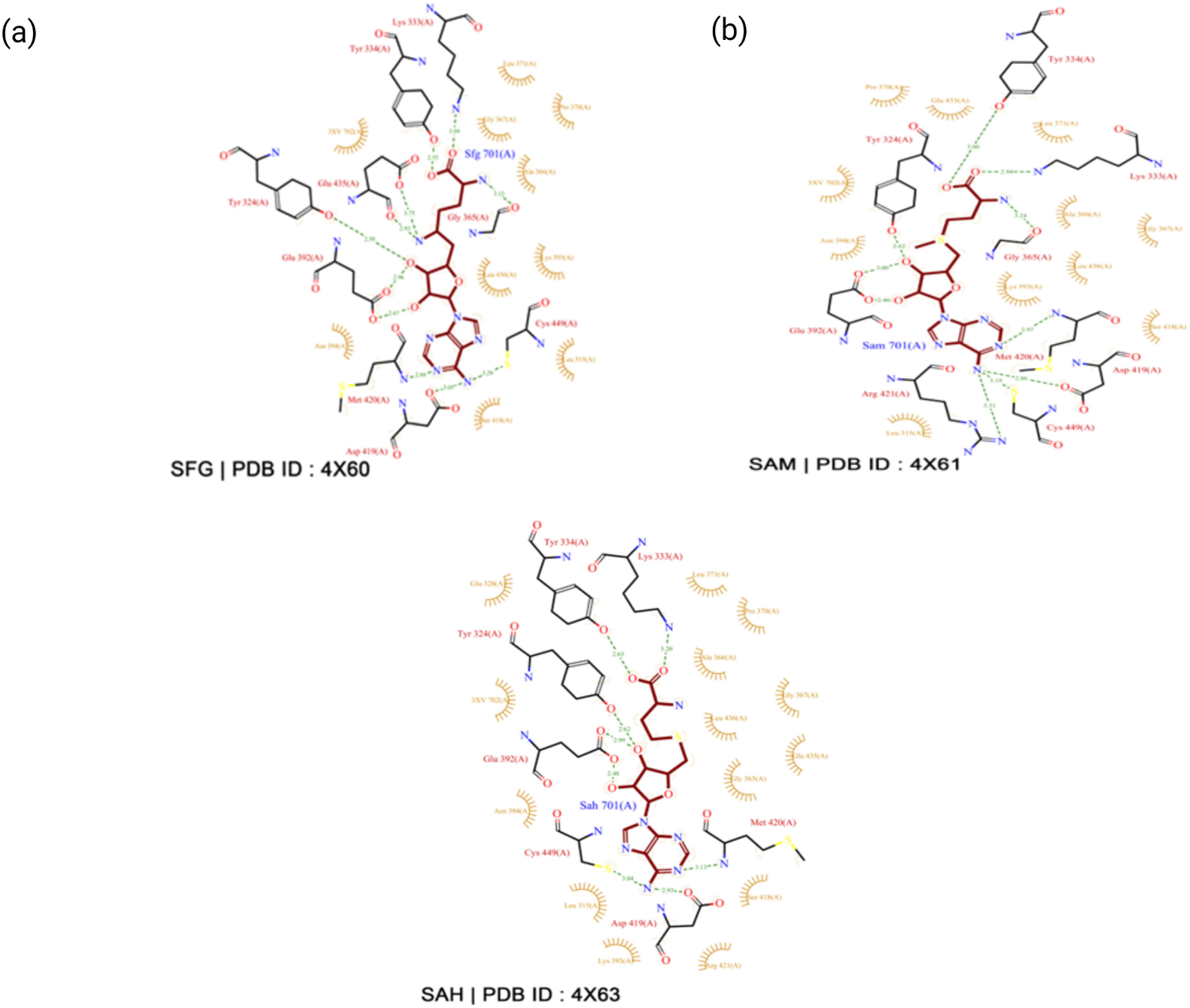
LigPlot interactions of EA with PRMT5 (PDB IDs: 4X60, 4X61, 4X63)

## Full blots images

**Figure-3:**
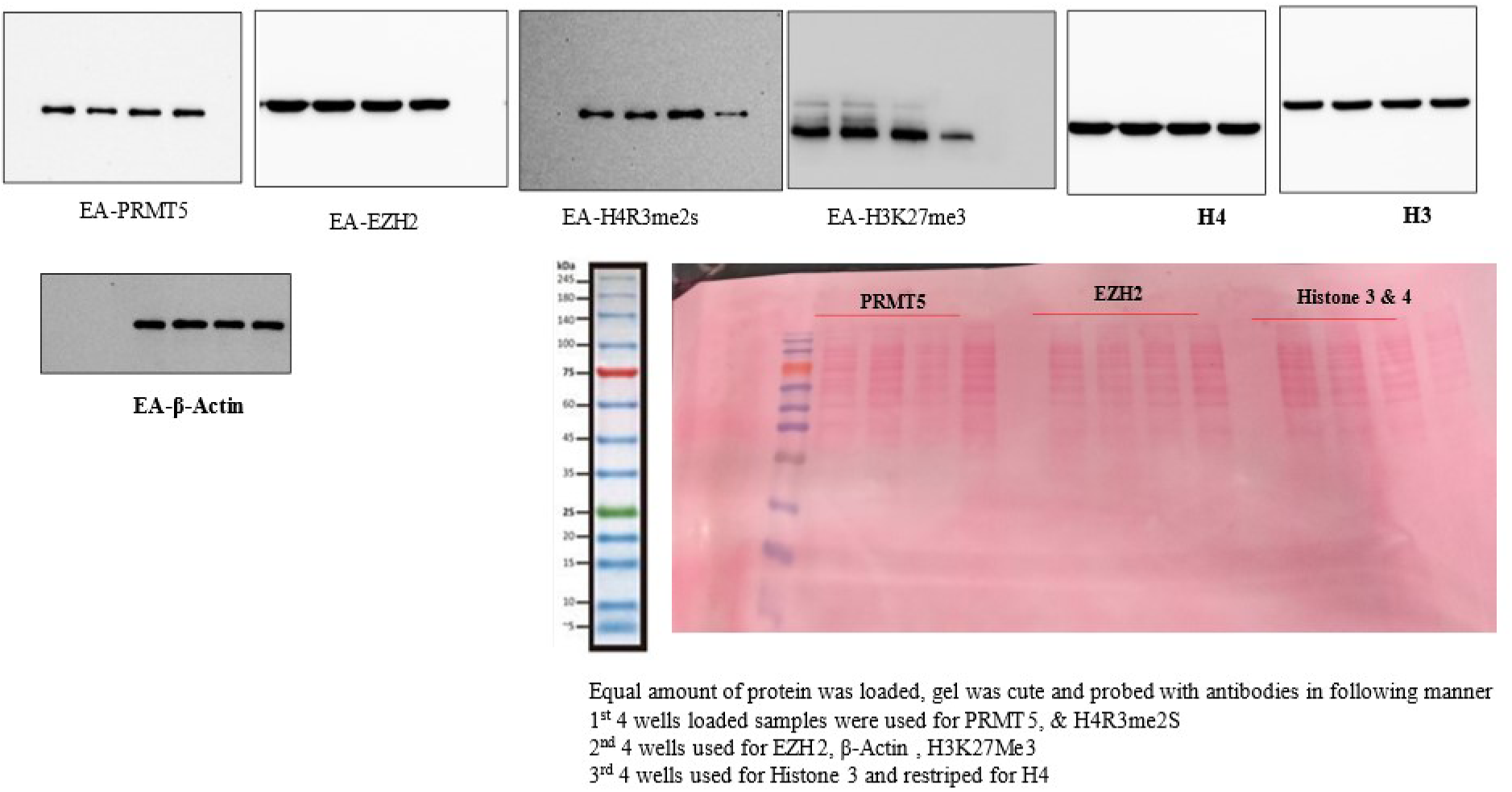
(Ellagic acid)

**Figure-6:**
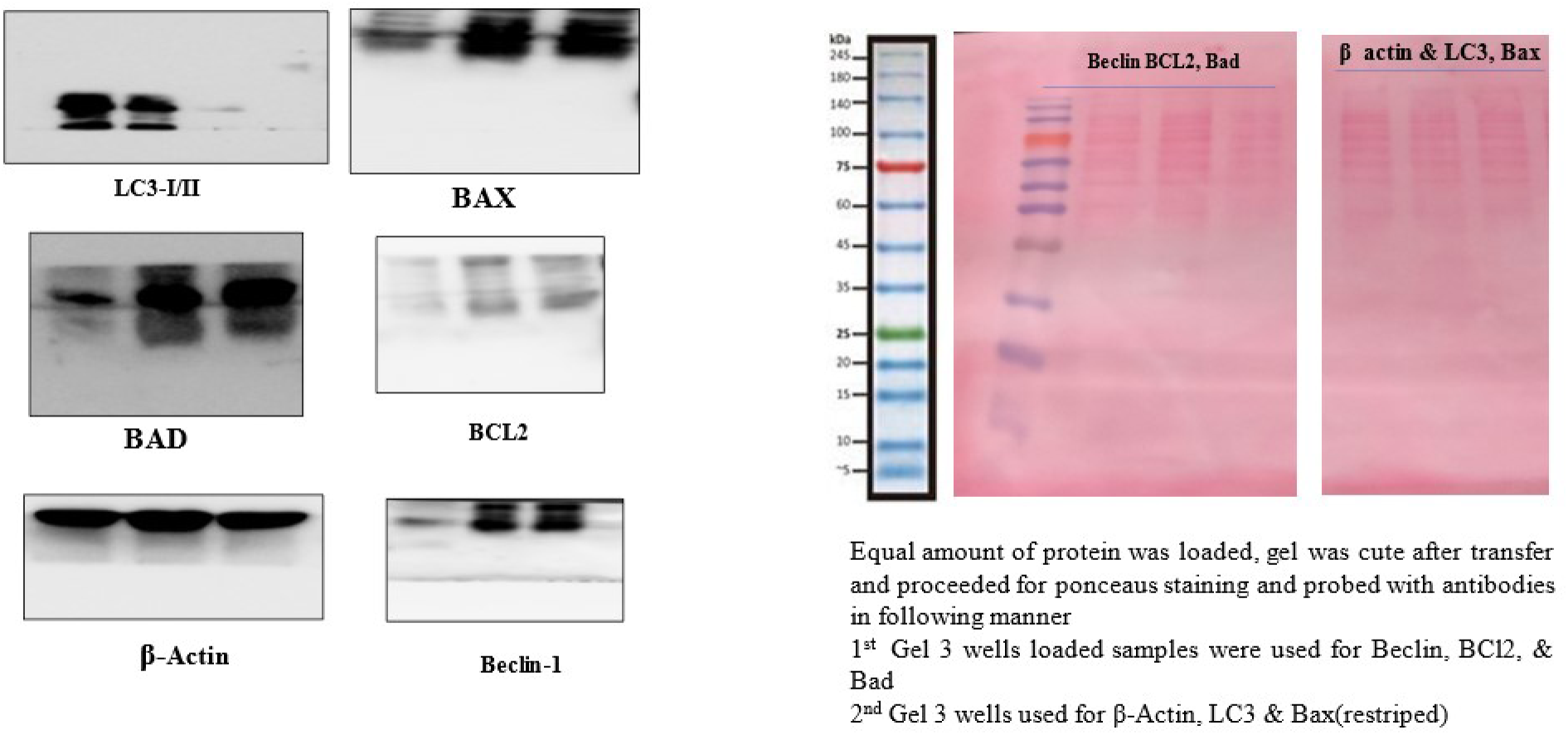
(EA blots)

**Figure-7:**
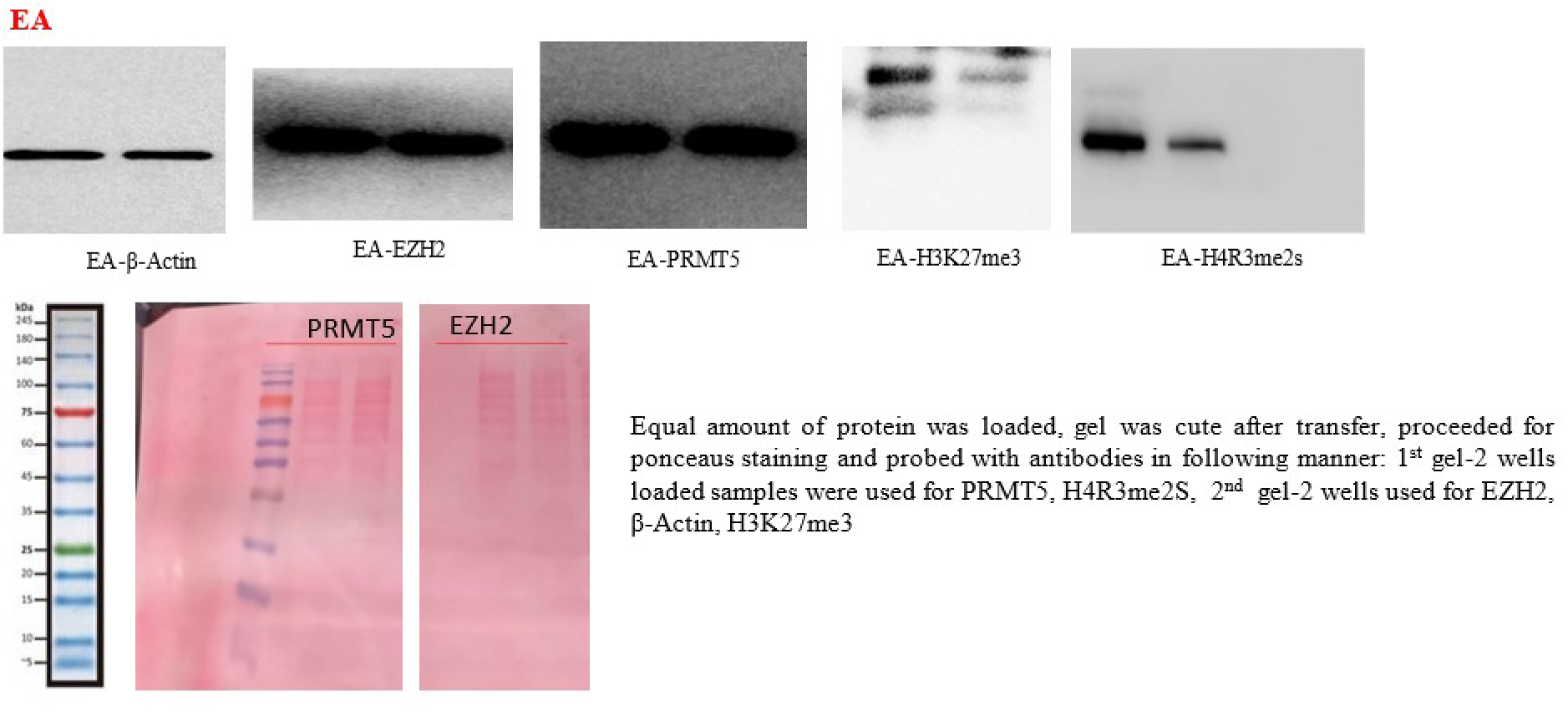
(EA blots)

